# Spatial grouping modulates the link between individual alpha frequency and temporal integration windows in crowding

**DOI:** 10.1101/2025.09.02.673743

**Authors:** Alessia Santoni, Luca Ronconi, Jason Samaha

**Affiliations:** Department of Psychology, Vita-Salute San Raffaele University, 20132 Milan, Italy; Department of Psychology and Cognitive Sciences, University of Trento, 38068 Trento, Italy; Department of Psychology, University of California, Santa Cruz, 95064 California, USA

## Abstract

Previous research has linked endogenous alpha oscillations (∼7–13 Hz) to temporal integration windows in visual perception, with higher individual alpha frequency (IAF) predicting improved temporal segregation. Here, we investigated whether alpha-rhythmic temporal integration is a factor in visual crowding and whether this relationship is mediated by spatial grouping mechanisms. 47 participants performed a Vernier discrimination task, in which we manipulated both the stimulus onset asynchrony (SOA) between flankers and targets, and the spatial configuration of the flankers. Specifically, flankers were arranged to either induce crowding or “uncrowding”, through the manipulation of good-Gestalt properties. Our results show that crowding has a temporal integration period of around 170 ms but this varies substantially across individuals. Importantly, resting-state IAF predicted individual variance in temporal integration windows: individuals with faster endogenous alpha rhythms could begin to segregate targets from distractors at shorter SOAs. Crucially, this effect was specific for crowding-inducing flankers and disappeared when flankers led to uncrowding. These results suggest that top-down spatial grouping can overwrite the temporal integration constraint imposed by alpha oscillations, highlighting both the relevance of alpha for understanding limits on peripheral visual processing as well as the flexible and context-dependent role of alpha in temporal integration.

**SIGNIFICANCE STATEMENT:** The frequency of alpha-band activity varies across individuals and has previously been linked to temporal integration windows for low-level stimulus properties (e.g., flashes). However, the relevance of individual alpha frequency (IAF) for everyday perception is less clear. We tested whether IAF predicts temporal integration in spatial crowding, a phenomenon that strongly limits perceptual abilities. We found that individuals with higher IAF were better able to spatially segregate targets from flankers at shorter flanker-target offsets, indicating that they spatially integrated the flankers over a smaller temporal window. Interestingly, this effect disappeared when crowding was alleviated by flanker configurations following good-Gestalt principles. Our results suggest that bottom-up mediated spatio-temporal constraints on perception are linked to alpha oscillations but can be overridden by top-down spatial cues.

## INTRODUCTION

When studying the visual system, we face a dual challenge: How does the brain parse sensory information in time, while simultaneously grouping elements of the scene across space? In other words, how does it construct perceptual objects that unfold coherently in both dimensions?

In the temporal domain, temporal integration refers to the idea that, although sensory input is continuous, the visual system does not process it in a seamless flow. Instead, perception might operate in a (quasi-)discrete or rhythmic fashion, where the timing of perceptual experience is defined by either perceptual ‘snapshots’ or ‘cycles’ (VanRullen, 2016; VanRullen & Koch, 2003), within which distinct stimulus features are integrated. The fact that both alpha oscillations (∼7–13 Hz) and the timing of perceptual processing across a wide range of psychophysical tasks share a similar cycle length (approximately 100 ms) has led to the idea that alpha activity may play a key role in temporal processing. In support of this view, a vast body of neurophysiological evidence has linked the temporal parsing of perceptual events around this temporal scale with the phase or frequency of alpha oscillations (e.g., Karvat & Landau, 2024; Kristofferson, 1967a, 1967b; Ronconi et al., 2017, 2024; Ronconi & Melcher, 2017; Samaha & Postle, 2015; Varela et al., 1981; Wutz et al., 2018). A common strategy to study temporal parsing is to use simple stimuli that flash isoretinally at varying temporal intervals: stimuli occurring closer in time are more likely to be perceived as a single, integrated percept. Notably, both between- and within-subject variations in the frequency of endogenous alpha rhythms have been linked to the temporal integration/segregation of visual input. More specifically, the resting-state Individual Alpha Frequency (IAF) has been previously found to predict subjective reports of simple flashes in a Two Flash Fusion Task (2FFT) (e.g., Deodato & Melcher, 2023; Drewes et al., 2022; Gray & Emmanouil, 2020; Haarlem et al., 2024; Samaha & Postle, 2015). Additionally, alpha frequency modulations can occur dynamically within individuals, based on task demands. For example, previous studies using a segregation-integration task where a target annulus was randomly positioned within a grid of distractors have shown that alpha frequency tends to increase or decrease when target identification relies on temporal segregation or integration mechanisms, respectively (Santoni, Di Dona, et al., 2025; Sharp et al., 2022; Wutz et al., 2018). Overall, converging evidence suggests that higher endogenous alpha frequencies are associated with enhanced temporal segregation mechanisms and, consequently, shorter temporal integration windows (for a review and meta-analysis, see Samaha & Romei, 2024).

Whether endogenous alpha rhythms coordinate both the temporal and spatial integration of visual information remains relatively unexplored. In the context of perceptual organization, uncrowding refers to an attenuation of crowding detrimental effects and occurs when flankers are based on good-Gestalt principles (Malania et al., 2007; Manassi et al., 2012, 2013; Morea et al., 2025; Santoni, Ronconi, et al., 2025; Sayim et al., 2014). This phenomenon reflects the visual system’s ability to reorganize cluttered input into coherent percepts, depending on the availability of meaningful grouping cues. The uncrowding effect has been previously interpreted in light of recurrent or feedback processing necessary to form a coherent Gestalt representation, as suggested by its relatively slow emergence (∼160 ms) under standard presentation conditions (Morea et al., 2025; Santoni, Ronconi, et al., 2025). Notably, previewing the flankers briefly before the full flanker-Vernier display (Morea et al., 2025), or presenting the flankers slightly earlier than the Vernier target (Santoni, Ronconi, et al., 2025), accelerates the uncrowding benefit (from 160 ms to 20-30 ms), supporting the idea that the early activation of grouping mechanisms can facilitate perceptual organization.

While temporal parsing and spatial grouping have typically been studied in isolation, natural vision requires their continuous interaction. Yet, little is known about whether temporal mechanisms, such as those linked to endogenous alpha oscillations, remain stable or adapt depending on the availability of spatial grouping cues. Given its temporal dynamics, uncrowding provides an ideal test case to investigate this question, as it reflects how spatial organization unfolds over time. Bridging this gap is critical for understanding how rhythmic neural activity supports the emergence of coherent percepts in dynamic environments.

In this study, we used a modified Vernier discrimination task in which we systematically varied the stimulus onset asynchrony (SOA) between Vernier targets and flankers to probe temporal integration and segregation. Crucially, we manipulated the spatial properties of the flankers so that they either induced crowding or uncrowding, allowing us to examine whether the link between IAF and temporal integration is modulated by the availability of grouping cues. Notably, we found that higher resting-state IAF promoted temporal segregation mechanisms selectively for the crowded condition, where spatial organization of Vernier-flankers configurations is disrupted. This finding suggests that endogenous alpha rhythms may impose limitations on temporal integration primarily through bottom-up mechanisms, but these limitations can be overridden in the face of top-down spatial cues.

## METHODS

### Participants

A total of 52 individuals from the University of California, Santa Cruz community took part in the study. Five were excluded due to poor performance (mean accuracy in the baseline condition < 0.6), yielding a final sample of 47 participants (average age = 20 years, SD = 3.14). Among the final group, 23 participants identified as female, 22 as male, and 2 as non-binary. Approximately 44% of participants identified as White/Caucasian, 13% as Asian, 17% as Hispanic/Latino, 2% as Middle Eastern, and 24% as mixed race. All participants had normal or corrected-to-normal vision, confirmed through a standard visual acuity assessment using a Snellen chart. The sample size was determined based on a recent meta-analysis examining the relationship between the IAF and temporal binding measures (Samaha & Romei, 2024), which indicated that a sample of 45 participants is needed to detect a population correlation of approximately 0.46 with 90% power using a two-tailed test. Participants received course credit for their participation, if desired. The research protocol was approved by the Institutional Review Board at the University of California, Santa Cruz.

### Apparatus and stimuli

Participants completed a Vernier discrimination task in a dimly lit room, seated 75 cm from the display with their head stabilized by a chinrest. Stimuli were rendered using Psychtoolbox (Brainard, 1997) for MATLAB (The MathWorks Inc., 2021) and displayed on a VIEWPixx/EEG monitor (120 Hz refresh rate) against a black background. Stimulus luminance was fixed at 80 cd/m².

Each Vernier stimulus was composed of two vertically aligned lines, with the lower line having a small horizontal offset. Each vertical line measured 40 arcminutes (′) in height and was spaced from the other by a vertical gap of 4′. The horizontal offset between the lines was set to 1′, with the direction (left or right) randomized across trials. Trials were separated by an intertrial interval of 1200 ms with a ±500 ms jitter. On each trial, the Vernier stimulus appeared randomly either to the left or right of a central fixation point, with an eccentricity of 4 degrees of visual angle from fixation. Both Vernier targets and flankers were presented for 40 ms. Flankers could be presented either simultaneously with the Vernier (0 ms SOA), before (SOAs: –166, –133, –100, –83, –66, –50, –33, –16 ms), or after the Vernier (33, 83, 166 ms), as shown in Figure 1B. The decision to include more negative SOAs was based on our primary interest in the effects of preceding flankers on target perception, which have previously been shown to elicit strong uncrowding effects even in the absence of temporal overlap (Santoni, Ronconi, et al., 2025).

**Figure 1.**
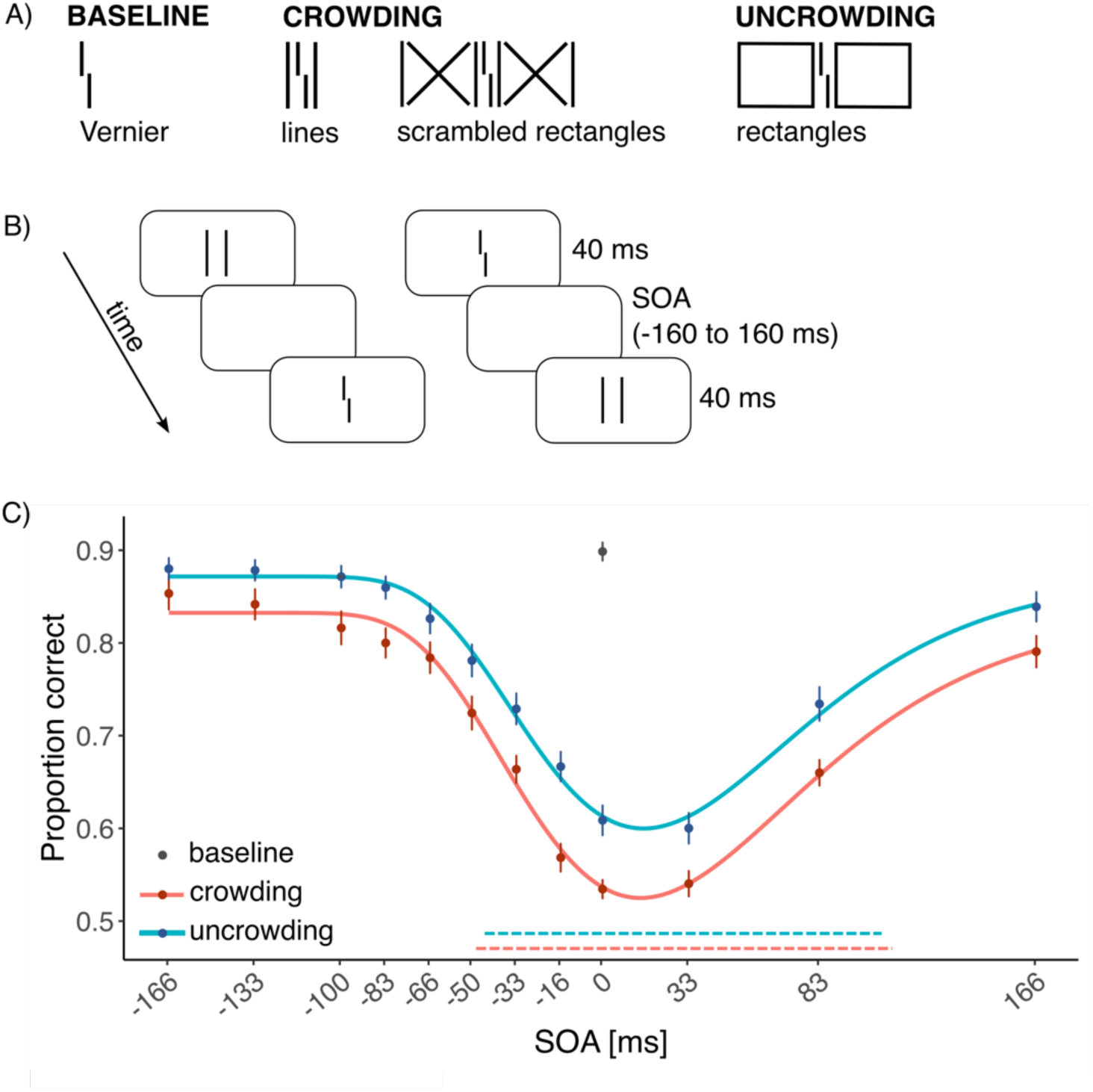
A) Stimuli used in the Vernier discrimination task. Flankers could be either lines/scrambled rectangles (leading to crowding), or rectangles (previously shown to elicit uncrowding). B) Flankers could be either simultaneous (SOA: 0 ms), precede (SOAs: –166, –133, –100, –83, –66, –50, –33, –16 ms), or follow (SOAs: 33, 83, 166 ms) Vernier targets. Participants were asked to report the horizontal offset of the lower bar of the Vernier stimulus. C) Behavioral performance is shown as colored dots (red = crowding, light blue = uncrowding, top dot at SOA 0 ms indicates performance in the baseline condition), with group-level inverted log-normal fits overlaid as lines. In the crowding condition, flankers were either vertical lines or scrambled rectangles; because both produced comparable effects, their data were combined and averaged. In the uncrowding condition, flankers were rectangular configurations that followed good-Gestalt principles. Performance shows an uncrowding benefit across stimulus onset asynchronies (SOAs) when good-Gestalt flankers are presented. Dashed horizontal lines at the bottom indicate group-level Temporal Integration Windows (TIWs) for crowding (red) and uncrowding (light blue) conditions, derived from the spread of the log-normal fits. Error bars represent the standard error of the mean (SEM).

Across different experimental blocks, the Vernier stimulus could either appear in isolation (baseline condition) or surrounded by flankers. Flankers could be either eliciting crowding or uncrowding, as previously demonstrated (Malania et al., 2007; Manassi et al., 2012, 2013; Morea et al., 2025; Santoni, Ronconi, et al., 2025; Sayim et al., 2014). In the uncrowding condition, rectangular flankers (44′ tall × 116′ wide) were arranged symmetrically around the target, at a distance of 16′. In the crowding condition, flankers were either simple vertical lines (44′ in height; 17 participants) or more complex scrambled rectangles (30 participants), selected to match, respectively, stimulus complexity at SOA 0 or the overall luminance of the uncrowding configuration (see Figure 1A).

### Procedure

Throughout the Vernier discrimination task, participants were instructed to maintain gaze on the fixation point, and report the Venier offset (i.e., offset of the lower line relative to the upper line) by pressing the “Z” key (left) or the “M” key (right) on a keyboard. Participants performed a total of 1016 trials: 56 baseline trials, 960 crowding and uncrowding trials (480 each). Baseline, crowding and uncrowding conditions were presented in separate blocks, counterbalanced across participants. Before the experiment, participants completed 20 practice trials with feedback to familiarize themselves with the procedure. They then performed the main task, which lasted approximately 50 minutes. EEG data were acquired both during task performance and during a subsequent eyes-closed resting-state period of 2 minutes. However, only the resting-state data were used here for analysis to maximize the detection of alpha activity in each subject. EEG signals were continuously recorded from 63 active electrodes using the BrainVision actiCHamp system (iMotions A/S, Copenhagen, Denmark). FCz served as the reference electrode during acquisition. Electrode impedances were kept below 10 kΩ, and data were sampled at 1000 Hz.

### Behavioral data analysis

First, we aimed to confirm the presence of the uncrowding effect for rectangular flankers as opposed to lines or scrambled rectangles. To this end, we conducted a repeated-measures ANOVA with Condition (2 levels: crowding vs. uncrowding) and SOA (12 levels: –166, –133, –100, –83, –66, –50, –33, –16, 0, 33, 83, 166 ms) as within-subject factors. Greenhouse-Geisser correction was applied when the assumption of sphericity was violated. Although the task included a greater number of negative SOAs to target the hypothesized uncrowding effect, the behavioral pattern showed evidence of uncrowding across both negative and positive SOAs (see Figure 1C). Therefore, to capture the full temporal profile, we fitted accuracy data using an inverted log-normal curve across the entire SOA range. This function captures the positively skewed, asymmetric shape previously observed in metacontrast masking tasks (e.g., (Bruchmann et al., 2010, 2016). The function was defined as follows:

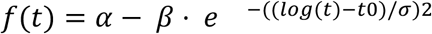

In this function, 𝑡 refers to the SOA (shifted to ensure all values are positive), α reflects the vertical offset, β scales the amplitude of the curve, t_0_ accounts for the location of the peak in log-time, σ controls the width or spread of the distribution. The expression t_0_ × (10^σ^−10^−σ^) was used to transform the parameter σ, originally defined in log space, back into milliseconds to capture the spread of the distribution in linear time, and was used as an estimate for the Temporal Integration Window (TIW). Parameters were estimated for each participant and condition using nonlinear least squares optimization via the *minpack.lm* package in R (Elzhov et al., 2022).

### EEG preprocessing and spectral analysis

EEG preprocessing was carried out using MATLAB (The MathWorks Inc., 2021), incorporating both custom-written scripts and functions from the EEGLAB toolbox (Delorme & Makeig, 2004). Resting-state EEG data were high-pass filtered at 0.1 Hz using the EEGLAB function *pop_eegfiltnew* and segmented into epochs of 1 second duration. The epoched data were resampled to 500 Hz and re-referenced using a median reference to prevent contamination from noisy channels during this stage of preprocessing. On average, 0.2% of the epochs (SD = 0.07) were rejected following visual inspection, and 0.48 channels (SD = 0.8) per participant were interpolated using spherical interpolation due to excessive noise. Subsequently, the data were re-referenced to the average reference.

The resting-state Individual Alpha Frequency (IAF) was computed from preprocessed resting-state EEG data using the *specparam* algorithm (Donoghue et al., 2020), via the FieldTrip toolbox (Oostenveld et al., 2011) in MATLAB. The algorithm isolates oscillatory activity from the aperiodic component of the power spectrum. Power spectral density was computed for each electrode via Fast Fourier Transform (FFT), applying a Hanning taper to each 1-second epoch. Data were zero-padded to achieve a frequency bin spacing of 0.125 Hz, allowing for finer estimation of spectral peak locations. The following parameters were used for fitting: minimum peak height = 0.001, maximum number of peaks = 3, peak threshold = 0.5, proximity threshold = 1 Hz, peak width limits = 0.5–12, and the aperiodic mode set to “fixed”. The spectral fits at ‘PO4’, the electrode with maximal alpha power, yielded a high overall quality (mean R² = 0.98, mean error = 0.005, R² range = 0.94–0.99).

### Behavior–IAF Analyses

To evaluate the relationship between TIWs and resting-state IAF, we performed two-tailed Spearman correlation analyses separately for each condition (crowding and uncrowding). For the correlation analyses, we considered the IAF value computed from the electrode showing maximal alpha power at the group level (electrode PO4), following previously established pipelines (e.g., Gray & Emmanouil, 2020; Ro, 2019; Samaha & Postle, 2015). Log-normal fits on behavioral data yielded a mean adjusted R² of 0.71 (SD = 0.20) for the crowding condition and a mean adjusted R² of 0.69 (SD = 0.21) for the uncrowding condition, reflecting reasonably consistent fit quality across participants. To ensure that TIW estimates were based on reliable fits, we restricted our main analyses to participants whose fits exceeded the average (adjusted R² > 0.7), thus including 29 participants in the crowding and 28 participants in the uncrowding condition. To ensure that our selection criterion did not introduce bias, we confirmed that adjusted R² values were not significantly correlated with TIWs (crowding: rho = 0.005, *p* = 0.98; uncrowding: rho = –0.29, *p* = 0.14). This step ensured that the a priori threshold did not bias TIW-based participant inclusion or artificially inflate IAF–TIW correlations due to uneven data quality.

In addition, we used a nonparametric cluster-based permutation approach implemented in FieldTrip to assess the spatial distribution of the correlation between IAF and TIWs across the whole scalp. Separately for each condition (crowding and uncrowding), we employed a cluster-level correction for multiple comparisons, which controls the family-wise error rate by evaluating the significance of clusters rather than individual channels (Maris & Oostenveld, 2007). We assessed cluster significance by comparing the observed Spearman correlation-based t-values against a null distribution built via the Monte Carlo method (number of random permutations = 2500) using a two-tailed test.

To test for the specificity of potential TIW-IAF effects, we run additional Spearman correlations between the IAF (at electrode PO4) and other fit parameters, namely: α, the vertical offset of the curve, β, the amplitude parameter, and t_0_, indicating the location of the peak. We also performed a Spearman correlation analysis between IAF and mean accuracy in the baseline condition (performed on all 47 participants), to verify that any observed effects were not simply due to general perceptual acuity in Vernier discrimination, rather than temporal integration/segregation mechanisms.

Finally, to further characterize the association between IAF and task performance in a model-free (i.e., without function fitting) analysis, we also fitted a linear mixed-effects model using the *lme4* package in R (Bates et al., 2015). The model predicted behavioral accuracy with the following fixed effects: Condition (crowding vs. uncrowding), resting-state IAF, SOA (entered as absolute values to account for potential nonlinear effects). Age was added as a covariate. All continuous predictors were z-scored prior to model fitting to facilitate interpretability and reduce multicollinearity, particularly in the presence of interaction terms. Participant was included as a random intercept to account for inter-individual variability. This approach enabled us to examine the contribution of IAF to task performance independently of the TIW fitting procedure, thereby including all participants in the sample (N=47).

## RESULTS

### Good-Gestalt flankers lead to uncrowding

To confirm the presence of the (un)crowding effects, we run a repeated-measures ANOVA on accuracy with Condition (crowding vs. uncrowding) and SOA (12 levels) as within-subject factors. A significant main effect of SOA, *F*(6.16, 283.4) = 178.20, *p* < .001 indicated that accuracy varied across temporal intervals between flankers and Vernier. A significant main effect of Condition also emerged, *F*(1, 46) = 55.18, *p* < .001, confirming overall higher accuracy in the uncrowding compared to the crowding condition (Figure 1C). The SOA × Condition interaction was not significant, *F*(6.56, 301.7) = 1.59, *p* = .14, suggesting that the magnitude of the uncrowding benefit did not differ substantially across SOAs. Additional analyses focusing on flanker type (i.e., scrambled rectangles, lines) within the crowding condition confirmed that both configurations reliably induced crowding. Notably, the SOA × Condition interaction reached significance in the subset of participants for whom the crowding-inducing flanker was a scrambled rectangle (see Figure S1 and Tables S1–S2 for detailed statistics by flanker configuration).

### IAF predicts TIWs under crowding conditions

As shown in Figure 2A, resting data from all 47 participants showed a clear peak in the power spectrum within the alpha range. Across the final sample, the average resting-state IAF was 10.3 Hz (SD = 0.72), consistent with previous findings in young adult populations (e.g., Ronconi et al., 2018; Samaha & Postle, 2015; Turner et al., 2023). Regarding the psychophysical measure, participants with good psychometric fit exhibited average TIWs of 169.62 ms (SD = 78) in the crowding condition (n = 29) and 133.33 ms (SD = 49.17) in the uncrowding condition (n = 28).

**Figure 2.**
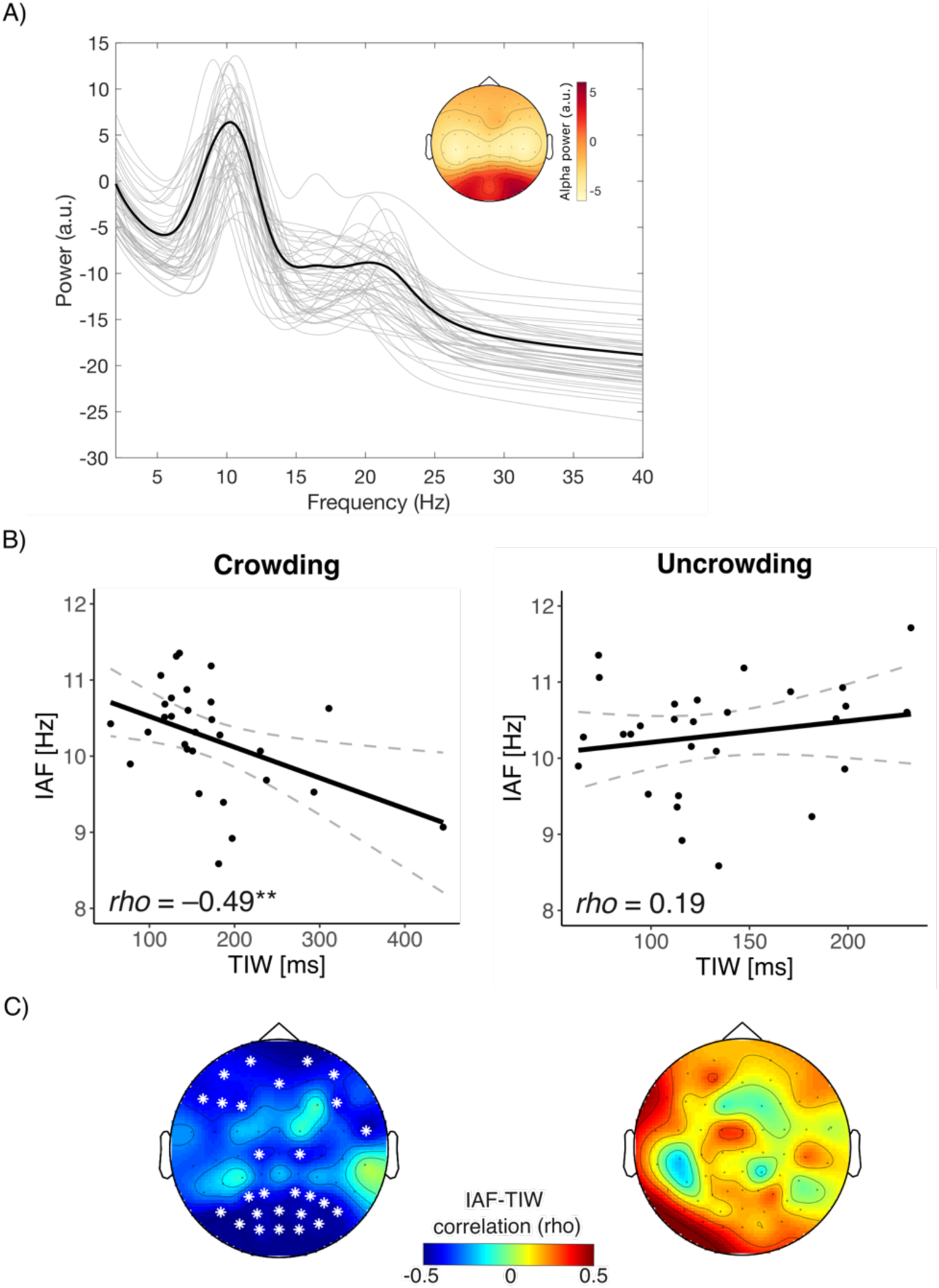
A) The modeled power spectrum at electrode PO4, estimated using the *specparam* algorithm, shows a prominent peak in the alpha range (7–13 Hz) for each participant (thin grey lines). The black line represents the average modeled spectrum across participants. In the upper right corner, the topographic distribution of alpha power, averaged across participants, is shown. B) A significant correlation emerged between Temporal Integration Windows (TIW) and the Individual Alpha Frequency (IAF) in the crowding condition (left panel; *rho* = –0.49, *p* = 0.008). Dots represent individual subjects; grey dashed lines represent 95% confidence intervals. C) This correlation was significant over parieto-occipital and frontal electrodes (left panel; *p* = 0.01, cluster corrected). No correlation between IAF and TIWs in the uncrowding condition emerged (B-C; right panels).

In support of our hypothesis, participants with higher resting-state IAF showed narrower TIWs in the Vernier discrimination task. Notably, this effect was specific to the crowding condition, as evidenced by a significant negative correlation (*rho* = –0.49, *p* = 0.008). In other words, individuals with faster alpha rhythms could temporally segregate flankers from Verniers and shorter SOAs, thereby mitigating crowding (Figure 2B, left panel). When exploring this correlation over the whole topography, a significant cluster emerged (*p_cluster_* = 0.01) over occipito-parietal and frontal electrodes (Figure 2C). In contrast, no significant correlation emerged in the uncrowding condition, neither when considering electrode ‘PO4’ (*rho* = 0.19, *p* = 0.33; Figure 2B, right panel) nor the whole topography of electrodes (Figure 2C, right panel).

To test if the observed pattern of results was specific to TIW and temporal segregation/integration mechanisms, we performed several control analyses. First, no significant correlation emerged between other fit parameters and IAF at PO4, indicating the specificity of the observed effect to TIW: α (crowding condition)-IAF: *rho* = 0.02, *p* = 0.9, α (uncrowding condition)-IAF: *rho* = –0.2, *p* = 0.93; β (crowding condition)-IAF: *rho* = –0.13, *p* = 0.5, β (uncrowding condition)-IAF: *rho* = –0.13, *p* = 0.52; t_0_ (crowding condition)-IAF: *rho* = –0.19, *p* = 0.32, t_0_ (uncrowding condition)-IAF: *rho* = –0.03, *p* = 0.87. Additionally, no correlation emerged between IAF and mean accuracy at the baseline condition, *rho* = –0.11, *p* = 0.47, suggesting the significant relationship with TIW was not driven by baseline differences in visual acuity.

Finally, we performed a complementary analysis that allowed us to consider the full final sample (N = 47) with no assumptions regarding the shape of the behavioral data (i.e., model-free). We fit a linear mixed-effects model predicting mean accuracy from Condition (crowding vs. uncrowding), resting-state IAF, and SOA, including their interactions, Age as a covariate, and a random intercept for participant. The model revealed a significant interaction between IAF and condition (β = –0.012, *SE* = 0.006, *t* = –1.99, *p* = 0.047), indicating that the relationship between IAF and performance differed across crowding conditions. Additionally, the model also revealed a main effect of Condition (β = 0.058, *p* < 0.001), IAF (β = 0.043, *p* < 0.021) and SOA (β = 0.122, *p* < 0.001), as well as an interaction between Condition and SOA (β = –0.014, *p* = 0.023; see Table S3 in Supplemental Materials for full model results).

## DISCUSSION

Since early work linking alpha rhythms to the temporal sampling of perceptual information (e.g., Kristofferson, 1967a, 1967b; Varela et al., 1981), neurophysiological research on this relationship has grown substantially. While most evidence supports a role for alpha frequency in temporal processing, fewer studies have questioned this link, reporting null results (e.g., Buergers & Noppeney, 2022; London et al., 2022; for reviews see Samaha & Romei, 2024; Schoffelen et al., 2024). We provide novel evidence that individuals with higher resting-state IAF exhibit narrower TIWs in visual crowding. To our knowledge, this is the first demonstration that alpha-frequency constrains temporal resolution in crowding. By contrast, in the uncrowding condition, where flankers adhered to good-Gestalt principles known to improve spatial grouping, no significant relationship between IAF and TIWs was observed. These findings suggest that spatial grouping mechanisms can override the fundamental limitation on temporal sampling imposed by alpha frequency.

Unlike other temporal integration paradigms where both displays are components of the target (e.g., two isoretinal flashes in the 2FFT or two complementary half-annuli in the segregation–integration task), the two consecutive displays in our study consisted of a target and a distractor that require *spatial* segregation for optimal performance. This key difference may explain why we observed a selective coupling between IAF and TIWs, which we propose reflects differences in the relative engagement of different neural mechanisms across the two conditions. In the uncrowding condition, spatial organization cues may have triggered perceptual grouping, at least partially relying on good-Gestalt principles. Previous evidence has shown that Gestalt cues, such as good continuation, may be resolved through a combination of feedforward and feedback processes, either within early visual areas (Marini & Marzi, 2016) or through recurrent interactions between early and higher-order visual regions (Wokke et al., 2013). While perceptual grouping cannot be solely attributed to Gestalt principles (Choung et al., 2023; Herzog, 2018), evidence has shown that similar feedforward and feedback interactions occur in (un)crowding (Chicherov & Herzog, 2015; Jastrzębowska et al., 2021; Ronconi et al., 2016; Ronconi & Bellacosa Marotti, 2017). This and other evidence have motivated theoretical accounts emphasizing the dynamic interplay between feedforward and feedback connections across the visual hierarchy (Lamme & Roelfsema, 2000; Roelfsema, 2006) and highlighting the limitations of purely feedforward or feedback models in explaining complex visual phenomena (Clarke et al., 2014).

Building on this evidence, we speculate that in the uncrowding condition, feedback connections may have activated the prototypical (i.e., good-Gestalt) flanker configurations, thereby reducing spatial interference and washing out the intrinsic temporal constraint imposed by alpha-rhythmic temporal integration. By contrast, when flankers violated grouping principles and provided no helpful spatial structure, crowding appeared to be primarily constrained by bottom-up limitations: target and distractors may initially be processed through a shared, capacity-limited channel early in the feedforward sweep, although later feedback could still play a modulatory role. Importantly, although both conditions might engage similar feedforward and feedback mechanisms, the selective IAF-TIW coupling reflects the relative weighting of these processes: when perceptual grouping is effective, feedback mitigates interference and temporal resolution is less dependent on alpha rhythms, whereas when grouping cues are absent, bottom-up processing dominates, making TIWs more directly linked to intrinsic alpha frequency. Together, these findings underscore the context-dependent role of alpha oscillations in shaping temporal integration during visual perception.

Our findings align with and extend prior work using Vernier discrimination tasks to study alpha oscillations. In this context, Menétrey et al. (2023) showed that Vernier discrimination deteriorates when a Vernier stimulus is followed by an anti-Vernier mask embedded within a stream of vertically aligned lines, with both alpha power and frequency modulating the reported offset. Similarly, Menétrey et al. (2024) used a backward masking paradigm and found that individuals with higher IAF were better able to segregate Vernier targets from temporally proximal masks, suggesting that alpha frequency influences temporal sampling across multiple cycles. Although these studies did not manipulate spatial organization, their findings reinforce our conclusion that alpha frequency supports temporal segregation when targets must be extracted from interfering input; a situation paralleled in our crowding condition.

Oscillatory contributions to spatio-temporal processing are not limited to alpha. In the spatial domain, stronger crowding conditions were linked to modulations of beta power alongside with alpha phase (Ronconi et al., 2016; Ronconi & Bellacosa Marotti, 2017). In the temporal domain, Ronconi et al. (2017, 2024), using both the 2FFT and apparent motion tasks, demonstrated that while decoding of 2FFT performance was strongest around alpha frequencies, perceptual decisions in the apparent motion task were linked to theta (6–7 Hz), consistent with its longer interstimulus intervals. From this perspective, the absence of an IAF-TIW relationship in uncrowding may reflect the recruitment of slower rhythms that support perceptual grouping. Behavioral work suggests that uncrowding emerges around a time window compatible with the alpha cycle when flankers are pre-activated, either by a preview (Morea et al., 2025) or temporal displacement (Santoni, Ronconi, et al., 2025). However, under standard presentation conditions, uncrowding unfolds slower, typical around 160 ms (Morea et al., 2025; Santoni, Ronconi, et al., 2025). Alternatively, the stronger engagement of feedback connections in the uncrowding condition might modulate alpha oscillations in a way that is not captured by resting-state IAF. Alpha rhythms are not a single, ubiquitous phenomenon but emerge from multiple cortical and subcortical generators (Bollimunta et al., 2008; Halgren et al., 2019) and only recently has work begun to disentangle these networks and their selective contributions to perception (e.g., Cruz et al., 2025; Nelli et al., 2021; Zhou et al., 2025). Identifying which alpha generators contribute to spatio-temporal sampling in different contexts remains an important goal for future research.

Additionally, the present results might have implications for neurodevelopmental conditions such as developmental dyslexia (DD), where both temporal processing deficits (Lallier et al., 2009; Ronconi et al., 2020; Santoni et al., 2024) and more pronounced crowding effects (Bertoni et al., 2019; Martelli et al., 2009; Spinelli et al., 2002) have been observed. Future work could test whether alpha-rhythms in temporal integration windows, previously shown to differ between typical and dyslexic readers (Santoni et al., 2025), could predict the severity of crowding-related reading impairments, and how these are modulated by perceptual grouping.

Taken together, our findings reveal that alpha oscillations predict temporal integration windows specifically under crowding, when bottom-up interference must be resolved in the absence of helpful grouping cues. By contrast, when Gestalt-based grouping can resolve spatial interference, the influence of alpha rhythms diminishes. This dissociation highlights the flexible and context-dependent role of alpha oscillations in shaping perceptual dynamics.

## ACKNOWLEDGMENTS

We would like to thank Emily Lincoln and Vrishab Nukala for their help in data collection. The present work was performed by A.S. in fulfilment of the requirements for obtaining the PhD degree at Vita-Salute San Raffaele University, Milan, Italy.

