## Supplemental Materials for "Spatial grouping modulates the link between individual alpha frequency and temporal integration windows in crowding"

**Figure S1.** Behavioral performance is plotted as a function of flanker configuration in the crowding condition. In the crowding condition, flankers could be either scrambled rectangles (N = 30) or single lines (N = 17), selected to match, respectively, the overall luminance of the uncrowding configuration or the stimulus complexity at baseline. Error bars represent the standard error of the mean (SEM).

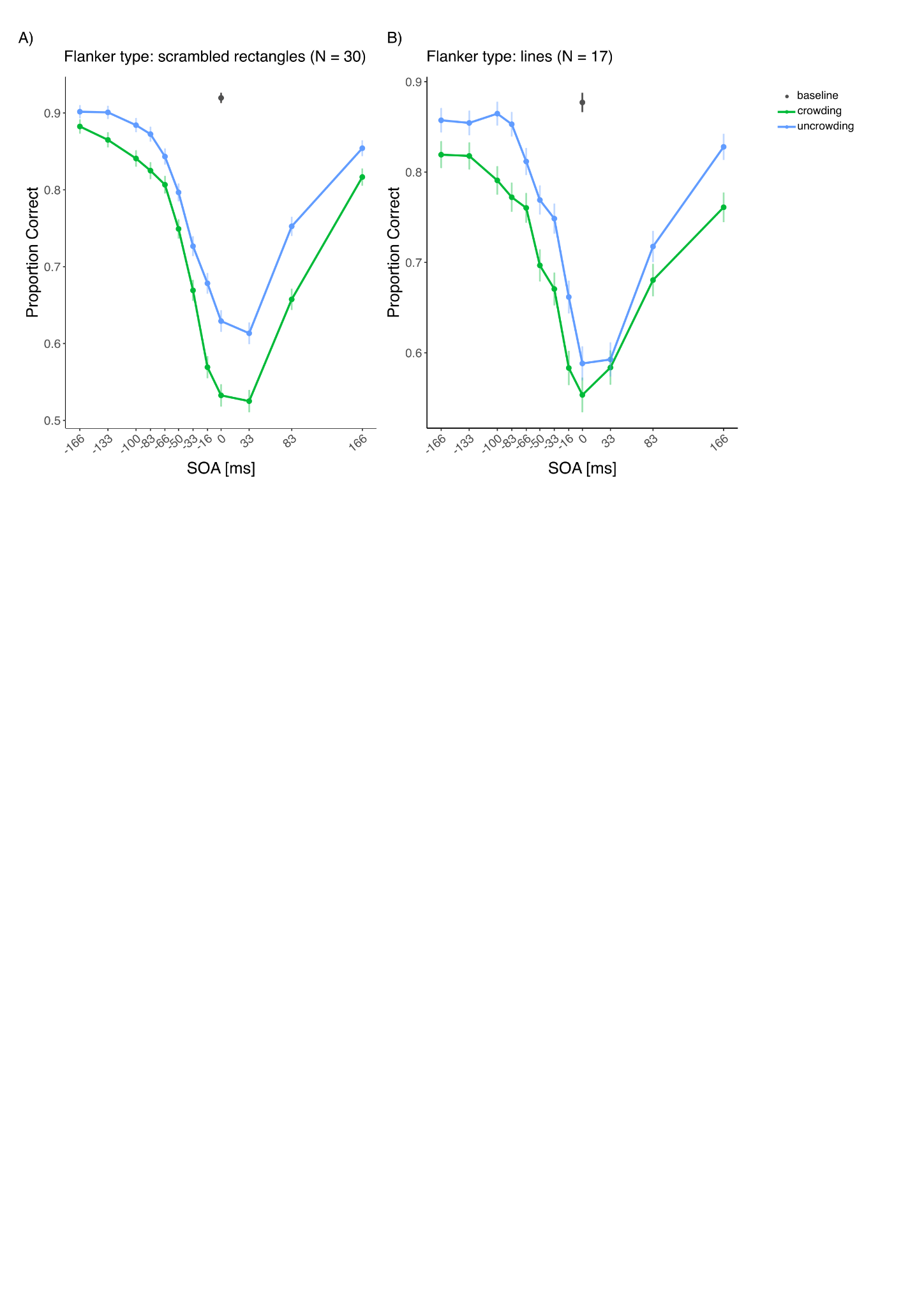

**Table S1.** Additional repeated-measures ANOVA results on accuracy data based on flanker configuration in the crowding condition. Both scrambled rectangles (N = 30) and simpler lines (N=17) led to crowding.

| **Effect** | **F (df)** | **p-value** |
| --- | --- | --- |
| **Flanker type: scrambled rectangles (N=30)** | | |
| SOA | 138.14 (7.59, 220.1) | < 0.001 |
| condition | 38.69 (1, 29) | < 0.001 |
| SOA × condition | 2.56 (6.29, 182.5) | 0.019 |
| **Flanker type: lines (N=17)** | | |
| SOA | 48.03 (5.23, 83.2) | < 0.001 |
| condition | 16.03 (1, 16) | 0.001 |
| SOA × condition | 0.85 (5.83, 92.9) | 0.529 |

**Table S2.** Post-hoc t tests for the SOA x condition interaction found when using scrambled rectangles as flankers in the crowding condition.

| **Contrast (crowding vs. uncrowding)** | **Estimate** | **SE** | **t (df)** | **p-value (FDR)** |
| --- | --- | --- | --- | --- |
| SOA: -166 vs. -166 | -0.0192 | 0.0182 | -1.055 (29) | 0.3001 |
| SOA: -133 vs. -133 | -0.0358 | 0.0141 | -2.548 (29) | 0.0224 |
| SOA: -100 vs. -100 | -0.0433 | 0.0171 | -2.538 (29) | 0.0224 |
| SOA: -83 vs. -83 | -0.0475 | 0.0171 | -2.772 (29) | 0.0189 |
| SOA: -66 vs. -66 | -0.0367 | 0.0182 | -2.014 (29) | 0.0582 |
| SOA: -50 vs. -50 | -0.0475 | 0.0149 | -3.190 (29) | 0.0102 |
| SOA: -33 vs. -33 | -0.0575 | 0.0212 | -2.715 (29) | 0.0189 |
| SOA: -16 vs. -16 | -0.1092 | 0.0252 | -4.339 (29) | 0.0009 |
| SOA: 0 vs. 0 | -0.0967 | 0.0219 | -4.421 (29) | 0.0009 |
| SOA: 33 vs. 33 | -0.0883 | 0.0287 | -3.079 (29) | 0.0108 |
| SOA: 83 vs. 83 | -0.0950 | 0.0228 | -4.163 (29) | 0.0010 |
| SOA: 166 vs. 166 | -0.0375 | 0.0175 | -2.140 (29) | 0.0491 |

**Table S3.** Results from a linear mixed-effects model (N=47) performed using the formula: mean_accuracy ~ condition * IAF * SOA (absolute value) + age + (1 | subject).

#### Random effects

| **Group** | **Effect** | **Variance** | **SD** |
| --- | --- | --- | --- |
| subject | Intercept | 0.00556 | 0.0746 |
| Residual |  | 0.01001 | 0.1000 |

#### Fixed effects

| **Predictor** | **Estimate** | **SE** | **df** | **t value** | **p value** |
| --- | --- | --- | --- | --- | --- |
| Intercept | 0.692 | 0.077 | 47.5 | 8.98 | < .001 |
| Condition | 0.058 | 0.006 | 1075 | 9.76 | < .001 |
| IAF | 0.043 | 0.019 | 292 | 2.32 | .021 |
| SOA | 0.122 | 0.015 | 1075 | 8.02 | < .001 |
| Age | –0.004 | 0.004 | 44 | –1.18 | .245 |
| Condition × IAF | –0.012 | 0.006 | 1075 | –1.99 | .047 |
| Condition × SOA | –0.014 | 0.006 | 1075 | –2.28 | .023 |
| IAF × SOA | 0.006 | 0.015 | 1075 | 0.39 | .697 |
| Condition × IAF × SOA | –0.004 | 0.006 | 1075 | –0.61 | .545 |
